# learnMET: an R package to apply machine learning methods for genomic prediction using multi-environment trial data

**DOI:** 10.1101/2021.12.13.472185

**Authors:** Cathy C. Westhues, Henner Simianer, Timothy M. Beissinger

**Author notes:** Corresponding authors:,. University of Goettingen, Department of Crop Sciences, Division of Plant Breeding Methodology, Center for Integrated Breeding Research, Carl-Sprengel-Weg 1, 37075 Goettingen, Germany.

## Abstract

We introduce the R-package *learnMET*, developed as a flexible framework to enable a collection of analyses on multi-environment trial (MET) breeding data with machine learning-based models. *learnMET* allows the combination of genomic information with environmental data such as climate and/or soil characteristics. Notably, the package offers the possibility of incorporating weather data from field weather stations, or can retrieve global meteorological datasets from a NASA database. Daily weather data can be aggregated over specific periods of time based on naive (for instance, non-overlapping 10-day windows) or phenological approaches. Different machine learning methods for genomic prediction are implemented, including gradient boosted trees, random forests, stacked ensemble models, and multi-layer perceptrons. These prediction models can be evaluated via a collection of cross-validation schemes that mimic typical scenarios encountered by plant breeders working with MET experimental data in a user-friendly way. The package is fully open source and accessible on GitHub.

## INTRODUCTION

Large amounts of data from various sources (phenotypic records from field trials, genomic or omics data, environmental information) are regularly gathered in the course of of multi-environment plant breeding trials (MET). The efficient exploitation of these extensive datasets has become of utmost interest for breeders to address essentially two objectives: (1) accurately predicting genotype performance in future environments; (2) untangling complex relationships between genetic markers, environmental covariables (ECs) and phenotypes to better understand the pervasive phenomenon of genotype-by-environment (G x E) interaction.

Several software packages have been recently developed to implement genomic prediction models that account for G x E effects, such as the following R packages: Bayesian Genomic Genotype × Environment Interaction (BGGE) (Granato *et al*. 2018), Bayesian Multi-Trait Multi-Environment for Genomic Selection (BMTME) (Montesinos-López *et al*. 2019) and EnvRtype (Costa-Neto *et al*. 2021b). These packages rely on Bayesian mixed models. BGGE presents a speed advantage explained by the use of an optimization procedure for sparse covariance matrices, while BMTME additionally exploits the genetic correlation among traits and environments to build linear G x E models. EnvRtype further widens the range of opportunities in Bayesian kernel models with the possibility to use non-linear arc-cosine kernels aiming at reproducing a deep learning approach (Cuevas *et al*. 2019; Costa-Neto *et al*. 2021a). This package also enables the retrieval of satellite-based weather data from the NASA POWER database.

While Bayesian approaches have been successful at dramatically improving predictive ability in multi-environment breeding experiments (Cuevas *et al*. 2017, 2019; Costa-Neto *et al*. 2021b), data-driven machine learning algorithms represent alternative predictive modeling techniques with increased flexibility with respect to the form of the mapping function between input and output variables. In particular, non-linear effects including gene x gene and gene x environment (G xE) interactions can be captured with machine learning models (Ritchie *et al*. 2003; McKinney *et al*. 2006; Crossa *et al*. 2019; Westhues *et al*. 2021). The latter are of utmost interest for plant breeders, as they can provoke a change in the relative ranking of genotypes across different environments, when cross-over interactions occur. Improved modeling of genotype-by-environment interactions through the inclusion of physiological stress indices that account for developmental stages, for instance obtained by the means of a crop growth model (Heslot *et al*. 2012; Rincent *et al*. 2017, 2019), can also be useful to leverage the full potential of machine learning in MET analyses.

Previous studies have shown that developing an all-purpose GP method is hardly possible due to various factors that affect the accuracy of the model. These include the training set size, the genetic architecture of the trait or the degree of relationship between training and test sets (Heslot *et al*. 2012; Bellot *et al*. 2018; Abdollahi-Arpanahi *et al*. 2020; Pook *et al*. 2020; Zingaretti *et al*. 2020). Therefore, a comparative analysis is often beneficial to identify the most suitable method for the intended use.

In this article we describe the R-package learnMET and its principal functionalities. learnMET provides an integrated pipeline to: (1) facilitate environmental characterization with several options for the user regarding the aggregation of daily weather data; and (2) evaluate various types of machine learning approaches with respect to their predictive performance based on relevant cross-validation schemes for MET datasets. The package offers flexibility by allowing to specify the sets of predictors to be used in predictions, or different methods to process genomic information to model genetic effects.

## METHODS

### Installation and dependencies

Using the devtools package (Wickham *et al*. 2021), learnMET can be easily be installed from GitHub and loaded (Box 1).

#### Box 1

**Install learnMET**

>devtools::install_github(“cjubin/learnMET”)

>library(learnMET)

Dependencies are automatically installed or updated when executing the command above.

### Real multi-environment trials datasets

Three toy datasets are included with the learnMET package to illustrate how input data should be provided by the user and how the different functionalities of the package can be utilized.

#### Rice datasets

The datasets were obtained from the INIA’s Rice Breeding Program (Uruguay) and were used in previous studies (Monteverde *et al*. 2018, 2019). Two breeding populations of rice (*indica*, composed of 327 elite breeding lines; and *japonica*, composed of 320 elite breeding lines) were phenotyped for four traits. The two populations were evaluated at a single location (Treinta y Tres, Uruguay) across multiple years and were genotyped using genotyping-by-sequencing (GBS) (Monteverde *et al*. 2019). Environmental covariables characterizing three developmental stages throughout the growing season were directly available.

#### Maize datasets

A subset of phenotypic and genotypic datasets collected and made available by the G2F initiative (www.genomes2fields.org) were integrated into learnMET. Hybrid genotypic data were obtained using GBS data from inbred parental lines. For more information about the original datasets, please refer to AlKhalifah *et al*. (2018) and McFarland *et al*. (2020). In total, phenotypic information collected from 22 environments covering 4 years and 6 different locations are included in the package.

### Running learnMET

learnMET is implemented as a three-step pipeline. These are described below.

#### Step 1: specifying input data and processing parameters

The first function in the learnMET pipeline is *create_METData* (Box 2). The user must provide genotypic and phenotypic data, as well as basic information about the field experiments (e.g. longitude, latitude, planting and harvest date). Climate covariables can be directly provided as day-interval aggregated variables, using the argument *climate_variables*. Alternatively, in order to compute weather-based covariables based on daily weather data, the user can set the *compute_climatic_ECs* argument to TRUE, and two possibilities are given. The first one is to provide raw daily weather data (with the *raw_weather_data* argument), which will undergo a quality control with the generation of an output file with flagged values, i.e. potential erroneous data. The second possibility, if the user does not have weather data available from measurements (e.g. from an in-field weather station), is the retrieval of daily weather records from the NASA POWER database using information contained in the *info_environments* argument, through the use of the package nasapower (Sparks 2018). Note that the function also checks which environments are characterized by in-field weather data in the *raw_weather_data* argument, in order to retrieve satellite-based weather data for the remaining environments without ground-based weather observations. An overview of the pipeline is provided in Figure 1.

##### Box 2

**Integration of input data in a METData list object**

***Case 1*: environmental covariables directly provided by the user**

>library(learnMET)

>data(geno_indica)

>data(map_indica)

>data(pheno_indica)

>data(info_environments_indica)

>data(env_data_indica)

>METdata_indica <create_METData(

geno = geno_indica,

map = map_indica,

pheno = pheno_indica,

climate_variables = climate_variables_indica,

info_environments = info_environments_indica,

compute_climatic_ECs = FALSE,

path_to_save = “/learnMET_analyses/indica”)

***Case 2*: daily climate data automatically retrieved and environmental covariables calculated via the package**

>data(geno_G2F)

>data(pheno_G2F)

>data(map_G2F)

>data(info_environments_G2F)

>METdata_g2f <-create_METData(

geno = geno_G2F,

pheno = pheno_G2F,

map = map_G2F,

climate_variables = NULL,

raw_weather_data = NULL,

compute_climatic_ECs = TRUE,

info_environments = info_environments_G2F,

soil_variables = soil_G2F,

path_to_save = “/learnMET_analyses/G2F”)

Note: code example to use ground-based daily weather data provided at https://cjubin.github.io/learnMET/articles/vignette_getweatherdata.html

**Figure 1.**
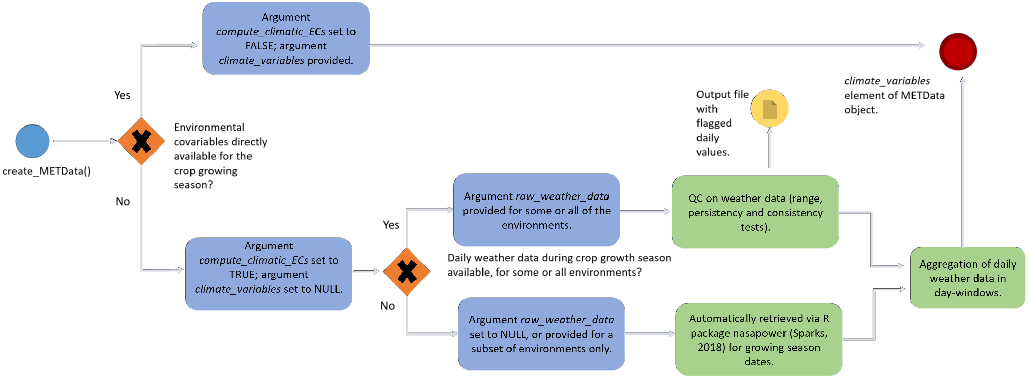
Overview of the pipeline regarding integration of weather data using the function *create_METData()* within the learnMET package. The blue circle signals the first step of the process, when the function is initially called. The blue boxes indicate how the arguments of the function should be characterized, according to the type of datasets available to the user. The green boxes indicate a task which is run in the pipeline via internal functions of the package. The red circle signals the final step, when the METData object is created. Details on the quality control tests implemented on daily weather data are provided at https://cjubin.github.io/learnMET/reference/qc_raw_weather_data.html, and on the methods to build environmental covariables based on aggregation of daily data at https://cjubin.github.io/learnMET/reference/get_ECs.html.

The aggregation of daily information into day-interval based values is also carried out within this function. Four methods are available and should be specified with the argument *method_ECs_intervals*: (1) default: use of a definite number of intervals across all environments (i.e. the duration of the windows vary according to the considered environment); (2) use of day-windows of fixed length (i.e. data are always aggregated over a fixed number of days, which can be adjusted by the user); (3) use of specific day intervals for each environment provided by the user, which should correspond to observed or assumed relevant phenological intervals; and (4) based on the estimated crop growth stage using accumulated growing degree-days in degrees Celsius.

Besides weather-based information, soil characterization for each environment can also be provided given the *soil_variables* argument. The output of *create_METData* is a list object of class *METData*, required as input for all other functionalities of the package.

#### Step 2: model evaluation through cross-validation

The second function in a typical workflow is *predict_trait_MET_cv()* (Box 3). The goal of this function is to assess a given prediction method using the specified cross-validation (CV) scenario. The CV schemes covered by the package are named according to the terminology employed by Jarquín *et al*. (2017) and correspond to real plant breeding prediction problems frequently evaluated using multi-environment datasets (Jarquín *et al*. 2014; Saint Pierre *et al*. 2016; Jarquín *et al*. 2017; Roorkiwal *et al*. 2018; Montesinos-López *et al*. 2018b,a; Costa-Neto *et al*. 2021a) and are defined as follows: (1) CV1: predicting the performance of newly developed genotypes (never tested in any of the environments included in the MET); (2) CV2: predicting the performance of genotypes that have been tested in some environments but not in others (also referred as field sparse testing); (3) CV0: predicting the performance of genotypes in new environment(s), i.e. the environment has not been tested; and (4) CV00: predicting the performance of newly developed genotypes in new environments, i.e. both environment and genotypes have not been observed in the training set. For CV0 and CV00, four configurations are implemented: leave-one-environment-out, leave-one-site-out, leave-one-year-out and forward prediction.

##### Box 3

**evaluation of a prediction method using a CV scheme**

>res_cv0_indica <-predict_trait_MET_cv(

METData = METdata_indica,

trait = “GC”,

prediction_method = “xgb_reg_1”,

cv_type = “cv0”,

cv0_type = “leave-one-year-out”,

compute_vip = TRUE,

seed = 100,

path_folder = “/user1/indica_cv_res/cv0”)

When *predict_trait_MET_cv()* is executed, a list of training/test splits is constructed according to the CV scheme chosen by the user. Each training set in each sub-element of this list is processed (e.g. standardization and removal of predictors with null variance, feature extraction based on principal component analysis), and the corresponding test set is processed using the same transformations. Performance metrics are computed on the test set, such as the Pearson correlation between predicted and observed phenotypic values (always calculated within the same environment, regardless of how the test sets are defined according to the different CV schemes), and the root mean square error. Analyses are fully reproducible given that seed and tuned hyperparameters are stored with the output of *predict_trait_MET_cv()*.

The function applies a nested CV to obtain an unbiased generalization performance estimate, implying an inner loop CV nested in an outer CV. The inner loop is used for model selection, i.e. hyperparameter tuning, while the outer loop provides the evaluation of model performance. Table 1 shows the different arguments that can be adjusted when executing the cross-validation evaluation.

**Table 1.** Description of the arguments used with the function predict_trait_MET_cv()

| Function argument | Description |
| --- | --- |
| METData | An object created by the initial function of the package create_METData(). |
| trait | Name of the trait to predict. |
| prediction_method | String to name the trait to predict. |
| lat_lon_included | Logical to use longitude and latitude as predictor variables. FALSE by default. |
| year_included | Logical to use year effect as dummy variable. FALSE by default. |
| cv_type | String indicating the cross-validation scheme to use among "cv0" (prediction of genotypes in new environments), "cv00" (prediction of new genotypes in new environments), "cv1" (prediction of new genotypes) or "cv2" (prediction of incomplete field trials). Default is "cv0". |
| cv0_type | String indicating the type of cv0 scenario, among "leave-one-environment-out", "leave-one-site-out", "leave-one-year-out" and "forward-prediction". Default is "leave-one-environment-out". |
| nb_folds_cv1 | Integer for the number of folds to use in the cv1 scheme, if it is selected. |
| repeats_cv1 | Integer for the number of repeats in the cv1 scheme, if it is selected.. |
| nb_folds_cv2 | Integer for the number of folds to use in the cv2 scheme, if it is selected. |
| repeats_cv2 | Integer for the number of repeats in the cv2 scheme, if it is selected.. |
| include_env_predictors | Logical to indicate if environmental covariables should be used in predictions. TRUE by default. |
| list_env_predictors | Vector of character strings with the names of the environmental predictors which should be used in predictions. NULL by default, which means that all environmental predictor variables are used. |
| seed | Integer with the seed value. Default is NULL, which implies that a random seed is generated and given as output. |
| save_processing | Logical to save the processing steps used to build the model in a RDS file. Default is FALSE. |
| path_folder | String to indicate the full path where the RDS file with results and plots generated during the analysis should be saved. |
| compute_vip | Logical to indicate whether the variable importance based each training set should be estimated. |
| num_pcs | Optional argument. Integer to indicate the number of principal components to derive from the genotype matrix or from the genomic relationship matrix (encouraged to speed up some analyses with cross-validation especially). |

#### Machine learning-based models implemented

Different machine learning-based regression methods are provided as S3 classes in an object oriented programming style. These methods are called within the integrated pipeline of the *predict_trait_MET_cv()* function.

In particular, the popular gradient boosting library XGBoost (Chen and Guestrin 2016), random forests (Breiman 2001), stacked ensemble models with Lasso regularization as meta-learners (Van der Laan *et al*. 2007), and multi-layer perceptrons using Keras (Chollet *et al*. 2015) are provided as prediction methods. Model stacking corresponds to an ensemble method which exploits the capabilities of many well-working models (called base learners) on a classification or regression task. For instance, support vector machines and gradient boosted tree models are trained as single learners on the same train-test splits, but can be sub-sampled for different sets of features and fitted using a grid of hyperparameter values, in an initial step. In the next step, a regularization method, like Lasso, learns how to combine their predictions to create a new model. Hence, the final model used for prediction only includes the best predictive models from the first step. Note that the classes we developed for pre-processing data and for fitting machine learning-based methods use functions from the tidymodels collection of R packages for machine learning (Kuhn and Wickham 2020), such as the Bayesian optimization to tune hyperparameters (function *tune_bayes()*) or stacked ensemble models (https://stacks.tidymodels.org/index.html). Feature importance can be estimated with model-agnostic methods (i.e. based on permutations) or with model-specific methods (e.g. gain metric for gradient boosted trees) using functions of the package DALEX (Biecek 2018) and vip (Greenwell *et al*. 2020).

#### Step 3: prediction of performance for untested genotypes and/or environments

The third module in the package aims at implementing predictions for unobserved configurations of genotypic and environmental predictors (Box 4). The user needs to provide a table of genotype IDs (e.g. name of new varieties) with their growing environments using the argument *pheno_new* in the function *add_new_METData()*. If new genotype IDs are included, corresponding genotypic data should be provided using the *geno_new* argument. Regarding characterization of new environments, the user can either provide a table of environments, with longitude, latitude and growing season dates, or can provide a table with their respective environmental covariables, in a similar manner as in the step 1. To build an appropriate model with learning parameters able to generalize well on new data, a hyperparameter optimization with cross-validation is conducted on the entire training dataset. Environmental covariables calculated for the test set in the *add_new_METData()* function should be provided or computed with the same data aggregation method (i.e. same *method_ECs_intervals*) as the one used for the training dataset in the step 1. Since it is exceedingly uncommon to have in-season weather data, one possibility is to use historical weather data at a location and obtain predictions across multiple years.

##### Box 4

**prediction of new observations using the complete training set**

>METdata_to_predict <-add_new_METData(

geno_new = geno_new,

METData_training = METdata_g2f,

pheno_new = pheno_new,

compute_climatic_ECs = TRUE,

info_environments_to_predict = info_environments_to_predict)

>new_envs_g2f <-predict_trait_MET(

METData_training = METdata_g2f,

METData_new = METdata_to_predict,

trait = “pltht”,

prediction_method = “xgb_reg_1”,

path_folder = “/user1/g2f/pred_new_environments”)

### Data Availability

The package is available on GitHub at https://github.com/cjubin/learnMET. Documentation and vignettes are provided at https://cjubin.github.io/learnMET/. Appendices with all scripts used to obtain the results presented in this paper can be found on GitHub at https://github.com/cjubin/learnMET/scripts_publication.

## RESULTS AND DISCUSSION

To illustrate the use of learnMET with multi-environment trials datasets, we provide here two example pipelines, both of which are available in the official package documentation. The first one demonstrates an implementation that requires no user-provided weather data, while the second pipeline shows prediction results obtained based on user-provided environmental data.

### Pipeline with maize data: retrieving meteorological data from NASA POWER database for each environment

When running the commands for step 1 (Box 1, Case 2) on the maize dataset, a set of weather-based variables (see documentation of the package) is automatically calculated using weather data retrieved from the NASA POWER database. By default, the method used to compute environmental covariables uses a fixed number of day-windows (10) that span the complete growing season within each environment. This optional argument can be modified via the argument *method_ECs_intervals* (detailed information about the different methods can be found at https://cjubin.github.io/learnMET/reference/get_ECs.html). The function *summary()* provides a quick overview of the elements stored and collected in this first step of the pipeline (Box 5).

#### Box 5

**Summary method for class METData**

> summary(METdata_g2f)

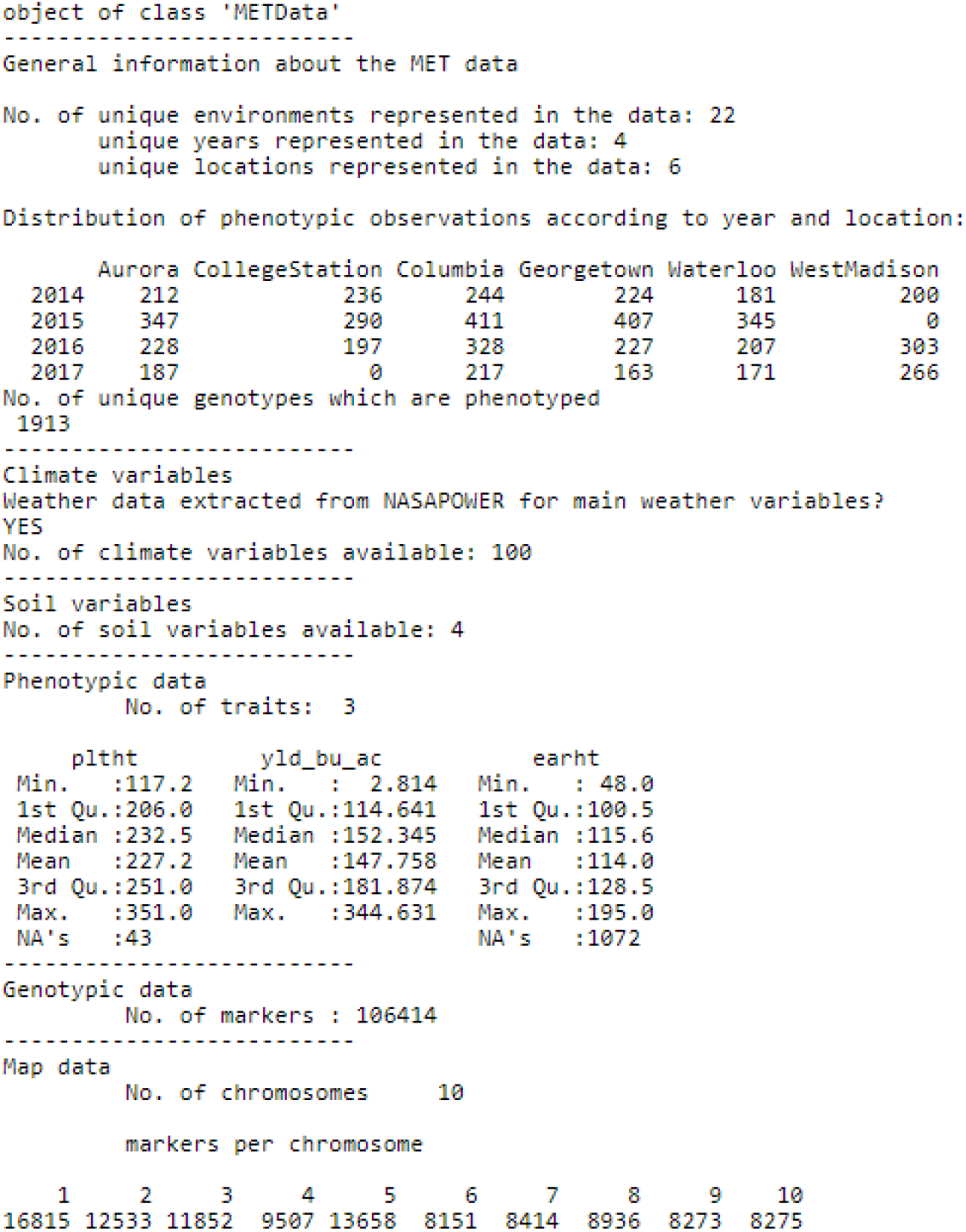

In addition, clustering analyses based on (a) only climate data; (b) only soil data (if available); and (c) all environmental variables together, are performed for a range of values for K = 2 to 10 clusters, and generated plots are saved on the path given by the argument *path_to_save* (Figure 2). Clustering analyses represent a useful tool to identify groups of environments with similar climatic conditions and to identify outliers, potentially associated with a low predictive ability in step 2.

**Figure 2.**
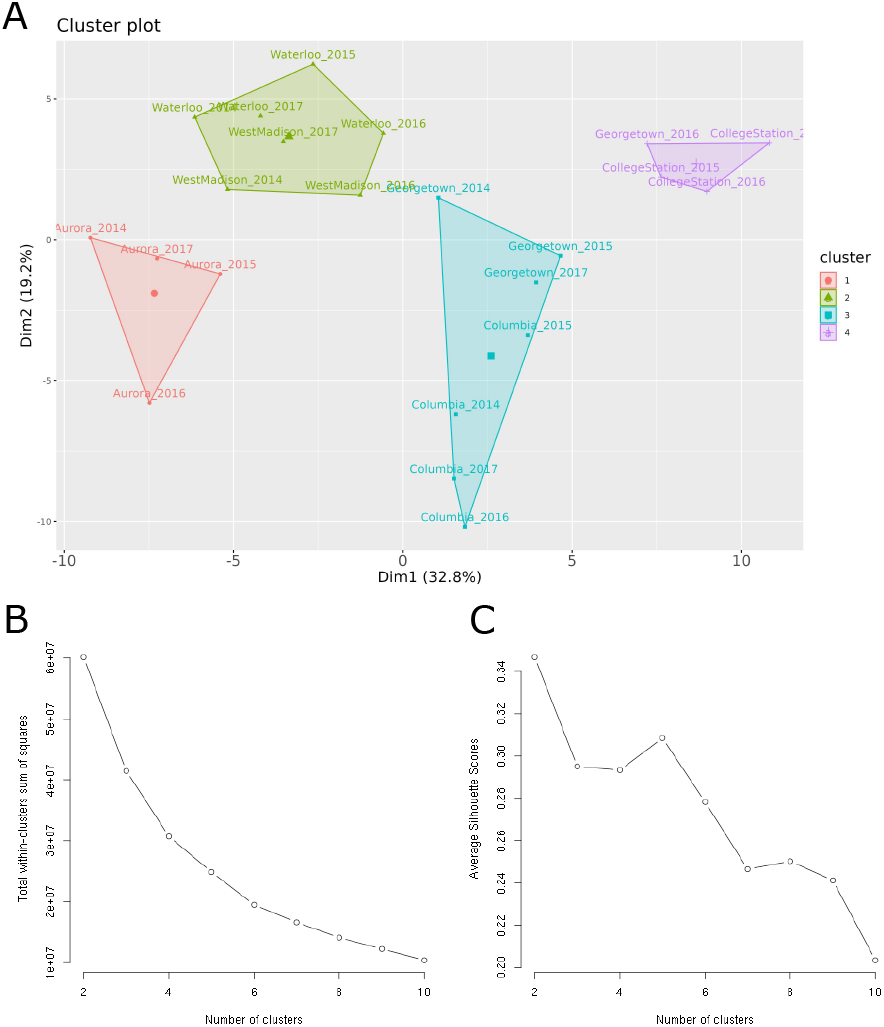
Output results from the *create_METData()* function. (A) Cluster analysis using K-means algorithm (K=4) to identify groups of similar environments based on climate and soil data. (B) Total within-cluster sum of squares as a function of the number of clusters. (C) Average Silhouette score as a function of the number of clusters. These methods can help users decide on the optimal number of clusters. Data used here is a subset of the Genomes to Fields maize dataset (AlKhalifah *et al*. 2018; McFarland *et al*. 2020). Weather data were retrieved from NASA POWER database via the package nasapower Sparks (2018).

### Pipeline with rice data: evaluation of two prediction methods with CV0-year

Cross-validation results from XGBoost and from stacked ensemble models on the rice toy dataset are presented in Table 2. The number of genomic predictors depends on the prediction method: in XGBoost, 100 principal components are used while in the stacked ensemble model, all SNPs are used. For XGBoost, Bayesian optimization is used to tune efficiently the number of boosting iterations, the learning rate and the depth of trees, generally considered as important hyperparameters controlling the risk of overfitting.

**Table 2.**
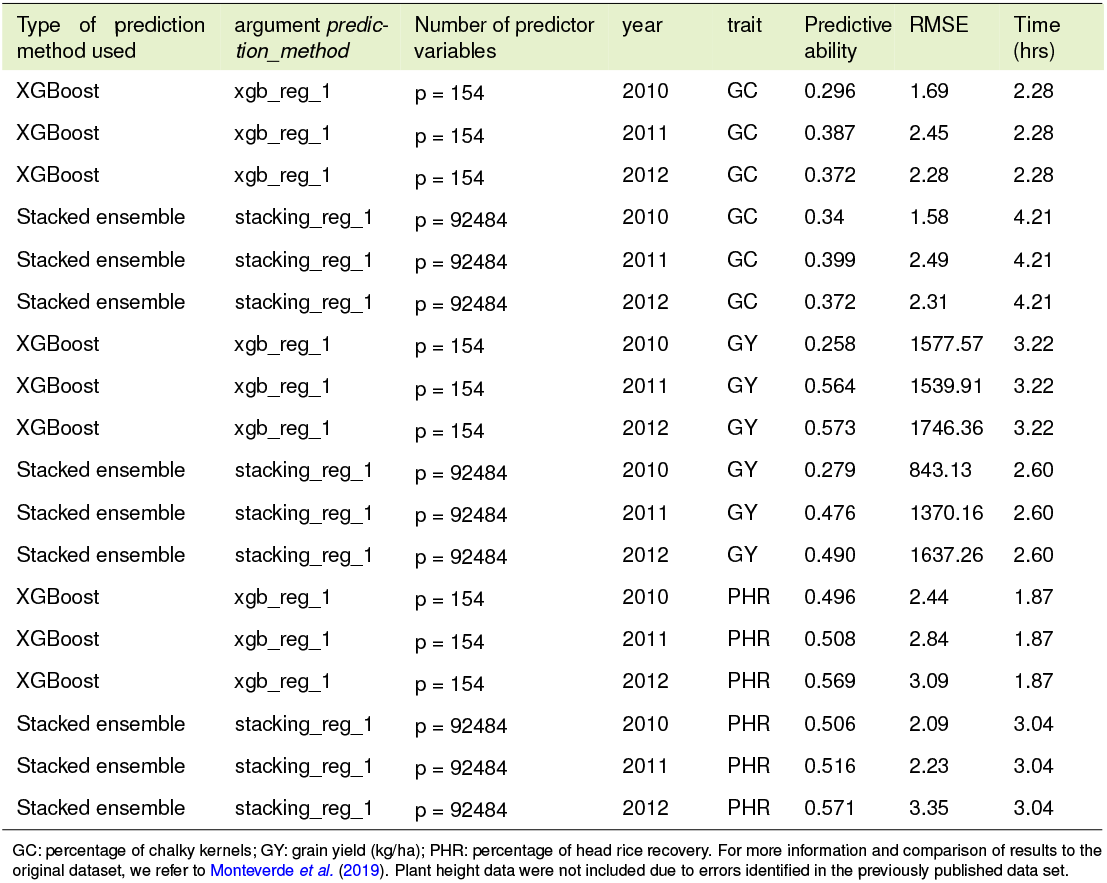
**Results from analyses conducted on the indica rice dataset (n = 981) for three phenotypic traits and two prediction methods using the function *predict_trait_MET_cv()*. A leave-one-year-out (CV0) cross-validation scheme was used. Predictive ability (estimated with the *r* Pearson correlation coefficient between predicted and observed values within the same environment), root mean square error (RMSE) and computation time for a complete trait-CV association are reported. The number of predictor variables depends on whether or not a dimensionality reduction method (e.g. principal component analysis) has been applied on the marker dataset. We used a node with 10 CPU cores, provided by the GWDG High Performance Computing Center of the University of Göttingen.**

| Type of prediction method used | argument *prediction_method* | Number of predictor variables | year | trait | Predictive ability | RMSE | Time (hrs) |
| --- | --- | --- | --- | --- | --- | --- | --- |
| XGBoost | xgb_reg_1 | p = 154 | 2010 | GC | 0.296 | 1.69 | 2.28 |
| XGBoost | xgb_reg_1 | p = 154 | 2011 | GC | 0.387 | 2.45 | 2.28 |
| XGBoost | xgb_reg_1 | p = 154 | 2012 | GC | 0.372 | 2.28 | 2.28 |
| Stacked ensemble | stacking_reg_1 | p = 92484 | 2010 | GC | 0.34 | 1.58 | 4.21 |
| Stacked ensemble | stacking_reg_1 | p = 92484 | 2011 | GC | 0.399 | 2.49 | 4.21 |
| Stacked ensemble | stacking_reg_1 | p = 92484 | 2012 | GC | 0.372 | 2.31 | 4.21 |
| XGBoost | xgb_reg_1 | p = 154 | 2010 | GY | 0.258 | 1577.57 | 3.22 |
| XGBoost | xgb_reg_1 | p = 154 | 2011 | GY | 0.564 | 1539.91 | 3.22 |
| XGBoost | xgb_reg_1 | p = 154 | 2012 | GY | 0.573 | 1746.36 | 3.22 |
| Stacked ensemble | stacking_reg_1 | p = 92484 | 2010 | GY | 0.279 | 843.13 | 2.60 |
| Stacked ensemble | stacking_reg_1 | p = 92484 | 2011 | GY | 0.476 | 1370.16 | 2.60 |
| Stacked ensemble | stacking_reg_1 | p = 92484 | 2012 | GY | 0.490 | 1637.26 | 2.60 |
| XGBoost | xgb_reg_1 | p = 154 | 2010 | PHR | 0.496 | 2.44 | 1.87 |
| XGBoost | xgb_reg_1 | p = 154 | 2011 | PHR | 0.508 | 2.84 | 1.87 |
| XGBoost | xgb_reg_1 | p = 154 | 2012 | PHR | 0.569 | 3.09 | 1.87 |
| Stacked ensemble | stacking_reg_1 | p = 92484 | 2010 | PHR | 0.506 | 2.09 | 3.04 |
| Stacked ensemble | stacking_reg_1 | p = 92484 | 2011 | PHR | 0.516 | 2.23 | 3.04 |
| Stacked ensemble | stacking_reg_1 | p = 92484 | 2012 | PHR | 0.571 | 3.35 | 3.04 |

GC: percentage of chalky kernels; GY: grain yield (kg/ha); PHR: percentage of head rice recovery. For more information and comparison of results to the original dataset, we refer to Monteverde et al. \(2019\). Plant height data were not included due to errors identified in the previously published data set.

Once a model has been evaluated with a CV scheme, various results can be extracted from the returned object, as shown in Box 6, and plots to visualize results are also saved in the *path_folder*.

#### Box 6

**Extraction of results from returned object of class *met_cv***

># Extract predictions for each test set in the CV scheme:

>pred_2010 <-res_cv0_indica$list_results_cv[[1]]$prediction_df

>pred_2011 <-res_cv0_indica$list_results_cv[[2]]$prediction_df

>pred_2012 <-res_cv0_indica$list_results_cv[[3]]$prediction_df

># The length of the list_results_cv sub-element is equal to the number of train/test sets partitions.

># Extract Pearson correlation between predicted and observed values for 2010:

>cor_2010 <-res_cv0_indica$list_results_cv[[1]]$cor_pred_obs

># Extract root mean square error between predicted and observed values for 2011:

>rmse_2011 <-res_cv0_indica$list_results_cv[[2]]$rmse_pred_obs

># Extract variable importance based on the model fitted to the training set composed of years 2010 and 2011:

>vip_2012 <-res_cv0_indica$list_results_cv[[3]]$vip

># Get the seed used:

>seed <-res_cv0_indica$seed_used

## CONCLUDING REMARKS AND FUTURE DEVELOPMENTS

*learnMET* was developed to make the integration of complex datasets originating from various data sources user-friendly. The package provides flexibility at various levels: (1) regarding the use of weather data, with the possibility to provide on-site weather stations data, or to retrieve external weather data, or a mix of both if on-site data are only partially available; (2) regarding how time intervals for aggregation of daily weather data are defined; (3) regarding predictive modeling: various types of nonlinear models are proposed, and we enable options to provide manually specific subsets of environmental variables (via the argument *list_env_predictors* in *predict_trait_MET_cv()*), and to specify which kernel functions to use in stacked ensemble models.

To allow analyses on larger datasets, future development of the package should include parallel processing to improve the scalability of the package and to best harness high performance computing resources. Improvements and extensions of deep learning models are also intended, as we did not investigate in-depth the network architecture (e.g. number of nodes per layer, type of activation function, type of optimizer) at this stage. Adding functions for feature engineering, such as filter methods to avoid keeping redundant predictors for prediction, could also potentially help improving both speed and predictive performance.

## ACKNOWLEDGMENTS

This work used the Scientific Compute Cluster at GWDG, the joint data center of Max Planck Society for the Advancement of Science (MPG) and University of Göttingen. We acknowledge support by the Open Access Publication Funds of the Göttingen University. The authors would like to thank the G2F Consortium for collecting data and making these publicly available. The authors are grateful to Eliana Monteverde for her useful input regarding the rice dataset, and also thank the National Institute of Agricultural Research (INIA-Uruguay) and technical staff from the experimental station from Treinta y Tres (Uruguay) for collecting the data.

## LITERATURE CITED

Abdollahi-Arpanahi, R., D. Gianola, and F. Peñagaricano, 2020 Deep learning versus parametric and ensemble methods for genomic prediction of complex phenotypes. Genetics Selection Evolution 52: 1–15.

AlKhalifah, N., D. A. Campbell, C. M. Falcon, J. M. Gardiner, N. D. Miller, et al., 2018 Maize genomes to fields: 2014 and 2015 field season genotype, phenotype, environment, and inbred ear image datasets. BMC Research Notes 11: 1–5.

Bellot, P., G. de Los Campos, and M. Pérez-Enciso, 2018 Can deep learning improve genomic prediction of complex human traits? Genetics 210: 809–819.

Biecek, P., 2018 Dalex: explainers for complex predictive models in r. The Journal of Machine Learning Research 19: 3245–3249.

Breiman, L., 2001 Random forests. Machine learning 45: 5–32.

Chen, T. and C. Guestrin, 2016 Xgboost: A scalable tree boosting system. In Proceedings of the 22nd acm sigkdd international conference on knowledge discovery and data mining, pp. 785–794.

Chollet, F. et al., 2015 Keras. https://keras.io.

Costa-Neto, G., R. Fritsche-Neto, and J. Crossa, 2021a Nonlinear kernels, dominance, and envirotyping data increase the accuracy of genome-based prediction in multi-environment trials. Heredity 126: 92–106.

Costa-Neto, G., G. Galli, H. F. Carvalho, J. Crossa, and R. Fritsche-Neto, 2021b Envrtype: a software to interplay enviromics and quantitative genomics in agriculture. G3 11: jkab040.

Crossa, J., J. W. Martini, D. Gianola, P. Pérez-Rodríguez, D. Jarquin, et al., 2019 Deep kernel and deep learning for genome-based prediction of single traits in multienvironment breeding trials. Frontiers in genetics 10: 1168.

Cuevas, J., J. Crossa, O. A. Montesinos-López, J. Burgueño, P. Pérez-Rodríguez, et al., 2017 Bayesian genomic prediction with genotype environment interaction kernel models. G3: Genes, Genomes, Genetics 7: 41–53.

Cuevas, J., O. Montesinos-López, P. Juliana, C. Guzmán, P. Pérez-Rodríguez, et al., 2019 Deep kernel for genomic and near infrared predictions in multi-environment breeding trials. G3: Genes, Genomes, Genetics 9: 2913–2924.

Granato, I., J. Cuevas, F. Luna-Vázquez, J. Crossa, O. Montesinos-López, et al., 2018 Bgge: a new package for genomic-enabled prediction incorporating genotype environment interaction models. G3: Genes, Genomes, Genetics 8: 3039–3047.

Greenwell, B. M., B. C. Boehmke, and B. Gray, 2020 Variable importance plots-an introduction to the vip package. R J. 12: 343.

Heslot, N., H.-P. Yang, M. E. Sorrells, and J.-L. Jannink, 2012 Genomic selection in plant breeding: a comparison of models. Crop science 52: 146–160.

Jarquín, D., J. Crossa, X. Lacaze, P. Du Cheyron, J. Daucourt, et al., 2014 A reaction norm model for genomic selection using highdimensional genomic and environmental data. Theoretical and applied genetics 127: 595–607.

Jarquín, D., C. L. da Silva, R. C. Gaynor, J. Poland, A. Fritz, et al., 2017 Increasing genomic-enabled prediction accuracy by modeling genotype x environment interactions in kansas wheat.

Kuhn, M. and H. Wickham, 2020 Tidymodels: a collection of packages for modeling and machine learning using tidyverse principles..

McFarland, B. A., N. AlKhalifah, M. Bohn, J. Bubert, E. S. Buckler, et al., 2020 Maize genomes to fields (g2f): 2014–2017 field seasons: genotype, phenotype, climatic, soil, and inbred ear image datasets. BMC research notes 13: 1–6.

McKinney, B. A., D. M. Reif, M. D. Ritchie, and J. H. Moore, 2006 Machine learning for detecting gene-gene interactions. Applied bioinformatics 5: 77–88.

Montesinos-López, A., O. A. Montesinos-López, D. Gianola, J. Crossa, and C. M. Hernández-Suárez, 2018a Multienvironment genomic prediction of plant traits using deep learners with dense architecture. G3: Genes, Genomes, Genetics 8: 3813–3828.

Montesinos-López, O. A., A. Montesinos-López, J. Crossa, D. Gianola, C. M. Hernández-Suárez, et al., 2018b Multi-trait, multienvironment deep learning modeling for genomic-enabled prediction of plant traits. G3: Genes, genomes, genetics 8: 3829–3840.

Montesinos-López, O. A., A. Montesinos-López, F. J. Luna-Vázquez, F. H. Toledo, P. Pérez-Rodríguez, et al., 2019 An r package for bayesian analysis of multi-environment and multitrait multi-environment data for genome-based prediction. G3: Genes, Genomes, Genetics 9: 1355–1369.

Monteverde, E., L. Gutierrez, P. Blanco, F. Pérez de Vida, J. E. Rosas, et al., 2019 Integrating molecular markers and environmental covariates to interpret genotype by environment interaction in rice (oryza sativa l.) grown in subtropical areas. G3: Genes, Genomes, Genetics 9: 1519–1531.

Monteverde, E., J. E. Rosas, P. Blanco, F. Pérez de Vida, V. Bonnecar-rère, et al., 2018 Multienvironment models increase prediction accuracy of complex traits in advanced breeding lines of rice. Crop Science 58: 1519–1530.

Pook, T., J. Freudenthal, A. Korte, and H. Simianer, 2020 Using local convolutional neural networks for genomic prediction. Frontiers in genetics 11: 1366.

Rincent, R., E. Kuhn, H. Monod, F.-X. Oury, M. Rousset, et al., 2017 Optimization of multi-environment trials for genomic selection based on crop models. Theoretical and Applied Genetics 130: 1735–1752.

Rincent, R., M. Malosetti, B. Ababaei, G. Touzy, A. Mini, et al., 2019 Using crop growth model stress covariates and ammi decomposition to better predict genotype-by-environment interactions. Theoretical and Applied Genetics 132: 3399–3411.

Ritchie, M. D., B. C. White, J. S. Parker, L. W. Hahn, and J. H. Moore, 2003 Optimizationof neural network architecture using genetic programming improvesdetection and modeling of gene-gene interactions in studies of humandiseases. BMC bioinformatics 4: 1–14.

Roorkiwal, M., D. Jarquin, M. K. Singh, P. M. Gaur, C. Bharadwaj, et al., 2018 Genomic-enabled prediction models using multienvironment trials to estimate the effect of genotype environment interaction on prediction accuracy in chickpea. Scientific reports 8: 1–11.

Saint Pierre, C., J. Burgueño, J. Crossa, G. F. Dávila, P. F. López, et al., 2016 Genomic prediction models for grain yield of spring bread wheat in diverse agro-ecological zones. Scientific reports 6: 1–11.

Sparks, A. H., 2018 nasapower: a nasa power global meteorology, surface solar energy and climatology data client for r.

Van der Laan, M. J., E. C. Polley, and A. E. Hubbard, 2007 Super learner. Statistical applications in genetics and molecular biology 6.

Westhues, C. C., G. S. Mahone, S. da Silva, P. Thorwarth, M. Schmidt, et al., 2021 Prediction of maize phenotypic traits with genomic and environmental predictors using gradient boosting frameworks. Frontiers in Plant Science 12: 2529.

Wickham, H., J. Hester, W. Chang, and M. J. Hester, 2021 Package ‘devtools’.

Zingaretti, L. M., S. A. Gezan, L. F. V. Ferrão, L. F. Osorio, A. Monfort, et al., 2020 Exploring deep learning for complex trait genomic prediction in polyploid outcrossing species. Frontiers in plant science 11: 25.

